# Fluorogenic Aptamer Optimizations on a Massively Parallel Sequencing Platform

**DOI:** 10.1101/2024.07.07.602435

**Authors:** Yu-An Kuo, Yuan-I Chen, Yanxing Wang, Zeynep Korkmaz, Siem Yonas, Yujie He, Trung D. Nguyen, Soonwoo Hong, Anh-Thu Nguyen, Sohyun Kim, Saeed Seifi, Po-Hsun Fan, Yuting Wu, Zhenglin Yang, Hung-Wen Liu, Yi Lu, Pengyu Ren, Hsin-Chih Yeh

## Abstract

<u>F</u>luorogenic <u>ap</u>tamers (FAPs) have become an increasingly important tool in cellular sensing and pathogen diagnostics. However, fine-tuning FAPs for enhanced performance remains challenging even with the structural details provided by X-ray crystallography. Here we present a novel approach to optimize a DNA-based FAP (D-FAP), Lettuce, on repurposed Illumina next-generation sequencing (NGS) chips. When substituting its cognate chromophore, DFHBI-1T, with TO1-biotin, Lettuce not only shows a red-shifted emission peak by 53 nm (from 505 to 558 nm), but also a 4-fold bulk fluorescence enhancement. After screening 8,821 Lettuce variants complexed with TO1-biotin, the C14T mutation is found to exhibit an improved apparent dissociated constant (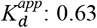 vs. 0.82 µM), an increased quantum yield (QY: 0.62 vs. 0.59) and an elongated fluorescence lifetime (τ: 6.00 vs. 5.77 ns), giving 45% more ensemble fluorescence than the canonical Lettuce/TO1-biotin complex. Molecular dynamic simulations further indicate that the π-π stacking interaction is key to determining the coordination structure of TO1-biotin in Lettuce. Our screening-and-simulation pipeline can effectively optimize FAPs without any prior structural knowledge of the canonical FAP/chromophore complexes, providing not only improved molecular probes for fluorescence sensing but also insights into aptamer-chromophore interactions.

---

Aptamers are single-stranded nucleic acids that bind specifically to a target ranging from small molecules^1^ to proteins^2^ and viral capsids^3^. Traditionally, aptamers are identified through systematic evolution of ligands by exponential enrichment (SELEX)^4-5^ from a pool of ∼10^15^ distinct strands and counter selected against structurally similar ligands to ensure recognition specificity. <u>F</u>luorogenic <u>ap</u>tamers (FAPs) are a special class of aptamers which light up upon binding their specific chromophore ligands^6^. RNA-based FAPs (R-FAPs) are especially useful for biological applications due to their genetically encodability^7-10^. Although endogenous probes based on R-FAPs have been demonstrated for cellular metabolite sensing in bacterial^10-11^ and mammalian cells^12-16^, the rapid degradation of RNA is a major issue of using R-FAPs in diverse applications.

Using SELEX, Jaffrey’s group identified a 100-nt-long DNA-based FAP (D-FAP) termed Lettuce that shows both enhanced chemical stability and improved photostability over its R-FAP counterpart Broccoli when complexing with the chromophore DFHBI-1T^17^. Whereas SELEX can identify a sequence that binds the designated chromophore most strongly and specifically within a pool of 10^15^ random strands, it is not guaranteed that the selected strand is the best FAP for the target chromophore. Post selection optimization of FAPs is needed and is often carried out on a small scale (<100 samples per experiment) with the aid of X-ray crystallography that provides the structural details of the FAP/chromophore complexes^18-20^. For instance, based on the crystal structure of the canonical Lettuce/DFHBI-1T complex, Ferré-D’Amaré’s group mutated the important chromophore interacting bases and unveiled the influence of these interacting bases on the photophysical properties of Lettuce/DFHBI-1T system^18^.

Although structure-guided optimization is feasible, it is a time-consuming approach to evolve an initially selected FAP. In addition, not all FAP/chromophore complexes have their crystal structures available (e.g., there was no published Broccoli/DFHBI structure). While SELEX can identify the best candidate among 10^15^ random sequences against a specific chromophore, there is a lack of effective methods to further scrutinize the sequence space (e.g., 10^4^) around the initial selection and continue to evolve the FAP design from there.

Illumina next-generation sequencing (NGS) chips contain millions to billions of sequenced polonies on the chip surface, making it an ideal platform to perform high-throughput mutagenesis study in protein and nanomaterial optimization^**21-34**^. By correlating the fluorescence of each polony to the sequence behind that spot, mutational impact on the fluorescence properties of diverse fluorescent molecules can be thoroughly investigated^24, 32^. Finkelstein’s group has previously established an NSG screening workflow called CHAMP (<u>c</u>hip-<u>h</u>ybridized <u>a</u>ssociation-<u>m</u>apping <u>p</u>latform) and used that to investigate ∼4,000 CRISPR-Cas complexes^26^. Using CHAMP, we also analyzed ∼40,000 fluorescent nanomaterials and identified the ones with unqiue emission properties^32^. Whereas Jaffrey’s group also showed the feasibility of visualizing R-FAPs on NGS chips, no detailed mutagenesis studies were performed in their report^35^.

Here we demonstrate an effective screening method termed D-FAP/CHAMP (<u>D</u>NA-based fluorogenic <u>ap</u>tamers on <u>c</u>hip-<u>h</u>ybridized <u>a</u>ssociation-<u>m</u>apping <u>p</u>latform) that can fluorescently characterize ∼10^4^ FAP sequence variants in a single experiment. D-FAP/CHAMP is made specifically for screening D-FAP/chromophore complexes and assessing the influence of mutations, insertions and deletions on complex’s photophysical properties in a massively parallel fashion.

Using D-FAP/CHAMP, here we screened a library of 8,821 distinct Lettuce variants, including single- and double-mutations, insertions, and deletions (**Figure 1**), against TO1-biotin and successfully identified the mutation C14T that gives 45% more ensemble fluorescence emission as compared to the canonical Lettuce. The reason that Lettuce was screened against TO1-biotin (M.W. = 749 g/mol, a chromophore typically used to light up the R-FAP Mango^36-37^) instead of the Lettuce’s cognate chromophore DFHBI-IT (M.W. = 320 g/mol) on CHAMP is because the resulting Lettuce/TO1-biotin complex showed an ensemble fluorescence emission intensity at least 4-fold more than that of the Lettuce/DFHBI-1T complex (under the same aptamer and chromophore concentrations using DPBS buffer; **Supplementary Figure S1C-D**). Moreover, the emission peak of Lettuce/TO1-biotin was found to be at 558 nm (ex: 526 nm), which is a 53 nm red shift from the 505 nm (ex: 455 nm) emission peak of Lettuce/DFHBI-1T complex. This red shift in emission greatly facilitated CHAMP screening. Our molecular dynamic (MD) simulations further demonstrated that TO1-biotin is bound to Lettuce and its variants primarily through the π-π stacking interactions, which is different from how DFHBI-IT is coordinated in Lettuce^18^. To the best of our knowledge, this is the first report that provides detailed mutagenesis studies on FAP optimization using repurposed Illumina NGS platform.

**Figure 1.**
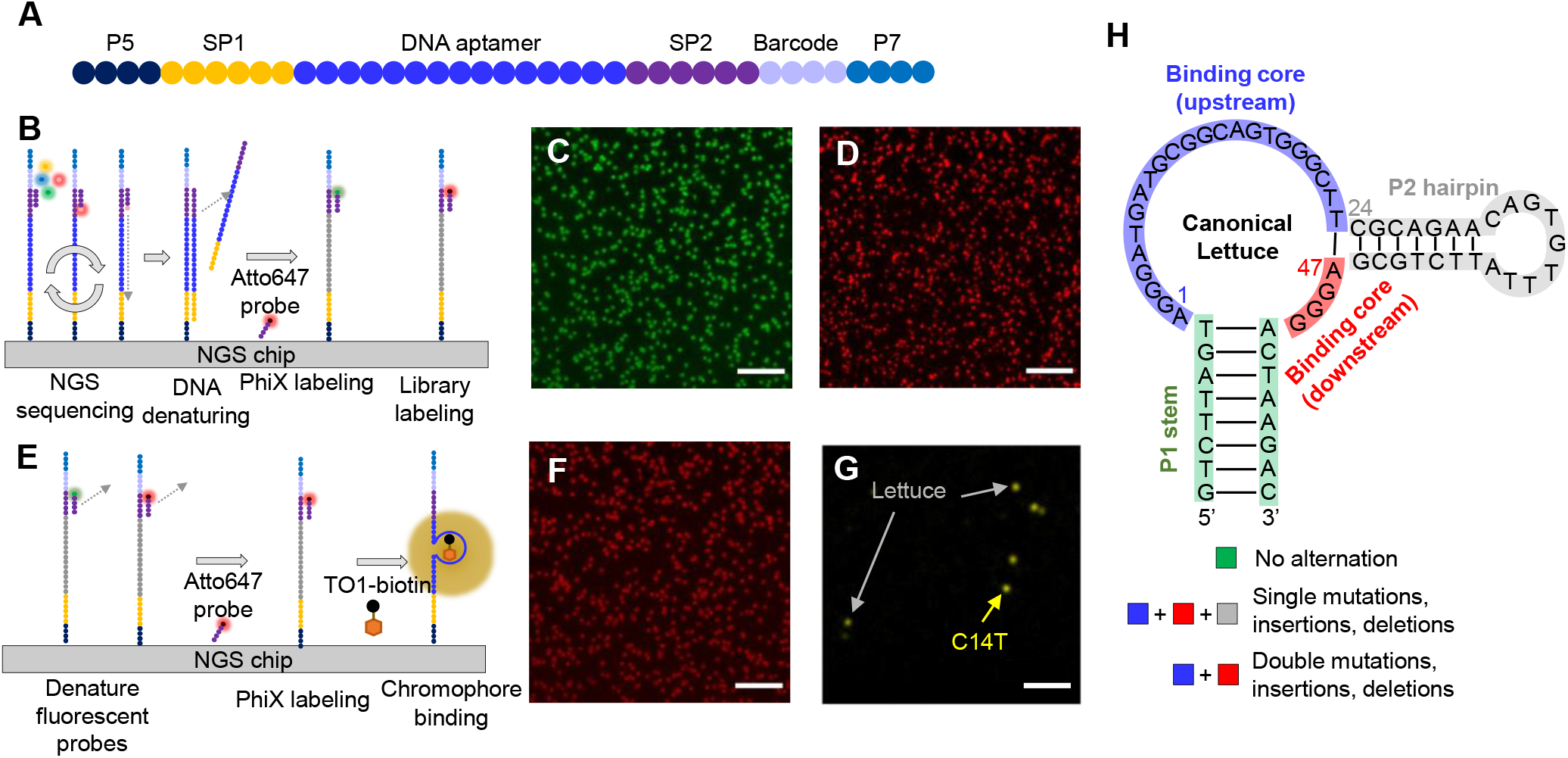
Overview of Lettuce aptamer optimization on Illumina NGS chips. (**A**) 8,821 distinct Lettuce variants are sandwiched in between Illumina sequencing adapters and synthesized on the flowcell surface. (**B**) NGS library was sequenced on a MiSeq chip through standard PE75 protocol and was denatured to expose ssDNA on the chip surface. Fiducial markers (i.e., PhiX) were labeled with Atto488N-tagged probe, followed by labeling library sequences with Atto647N-tagged probe. The chip was then visualized on an epi-fluorescence microscope to evaluate the polony density and chip quality. (**C**) Representative images of PhiX labeling and (**D**) library labeling, respectively. (**E**) To image TO1-biotin on the chip surface, we first denatured the labeling probes in (**B**) and label PhiX with Atto647N-tagged probes, followed by incubating flowcell with 200 nM TO1-biotin. (**F**) A representative image of PhiX image labeled with Atto647N-tagged probes. (**G**) A zoom-in image of the light-up variants after CHAMP alignment. (**H**) Schematic of canonical Lettuce and library design in this study. Scale bars in (**C-D**) and (**F**) are 20 µm, while scale bar in (**G**) is 10 µm.

Please note that the Lettuce sequence reported in the follow-up article by Ferré-D’Amaré’s group featured a shortened P1 stem and P2 hairpin (hereafter denoted as sLettuce and sC14T, **Supplementary Figure S1A-B**)^18^ as compared to the original Lettuce sequence^17^. However, the chromophore binding core remained unchanged. In this report, if not otherwise stated, Lettuce refers to the FAP sequence published in the Jaffrey group report^17^.

The D-FAP/CHAMP screening workflow started with adding adapters and barcodes to the library design (**Figure 1A**). The full-scale library strands were then captured by surface adapters, sequenced (**Figure 1B**) and imaged separately with a PhiX fiducial marker label (i.e., an Atto488N-tagged probe, **Figure 1C**) and a library label (i.e., an Atto647N-tagged probe, **Figure 1D**) on an Illumina *MiSeq* chip (**Experimental Section** in **Supplementary Information**). The two labels were then denatured, and solutions of 500 µM red fiducial marker label and 200 nM TO1-biotin were introduced to the chip sequentially (**Figure 1E**). The labeled fiducial marker and the light-up Lettuce variants were imaged with two different filter cubes (a standard Cy5 filter cube (Ex/Em: 620/60 nm, 700/75 nm) for the fiducial markers and a modified FITC filter cube (Ex/Em: 480/40 nm, 560/40 nm) for the light-up Lettuce). An image alignment algorithm and a postprocessing pipeline developed in house^26, 32^ was then used to identify the aptamer sequence behind each light-up polony (**Figure 1G**). On average each aptamer design had 95 duplicated polonies on the chip surface (with a standard deviation of 116), thus giving a highly reliable and reproducible screening result.

Our CHAMP screening results were summarized as the fluorescence improvement ratio of each variant in the heatmaps (**Figure 2A** and **Supplementary Figure S3-S5**). An improvement ratio of zero represented no improvement when compared to the canonical Lettuce/TO1-biotin complex, while improvement indices of +19.1 (C14T) and -5.7 (A5T) represented a 19.1% increase and a 5.7% decrease in the CHAMP observed fluorescence, respectively. From the single-mutation analysis, it was clear to see that A5, T9, C14, T22 and G51 bases play a significant role in controlling the photophysical property of Lettuce/TO1-biotin complex, as some mutations at these locations can give an improvement ratio greater than -10% (e.g., -7.5% at T9A). Our observation well agreed with the crystal structures published by Ferré-D’Amaré’s group, where these five bases were indeed in close proximity to the encapsulated chromophore and thus sensitive to the mutations^18^ (**Figure 2B**). Our screening results also indicated that the sequences in the Q1 and Q2 tiers of the G-quadruplex need to be conserved – any destabilization of the G-quadruplex deteriorates the chromophore binding core, leading to a decrease in fluorescence emission. In contrast, mutations in the P2 hairpin loop (C30-A39 in **Figure 2C** and **Supplementary figure S3-S5**) did not affect the fluorescence emission of Lettuce/TO1-biotin, suggesting the loop bases are not interacting with the chromophore. But the P2 hairpin stem (C24-A27 and T43-G46) was found mutation intolerant as this stem portion is still close to the encapsulated chromophore according to the crystal structure^18^. Please note that in this report the starting position (A1) is the first nucleobase of the Lettuce binding core, which is different from the earlier report where the starting position is at the 5’ end of the stem^18^.

**Figure 2.**
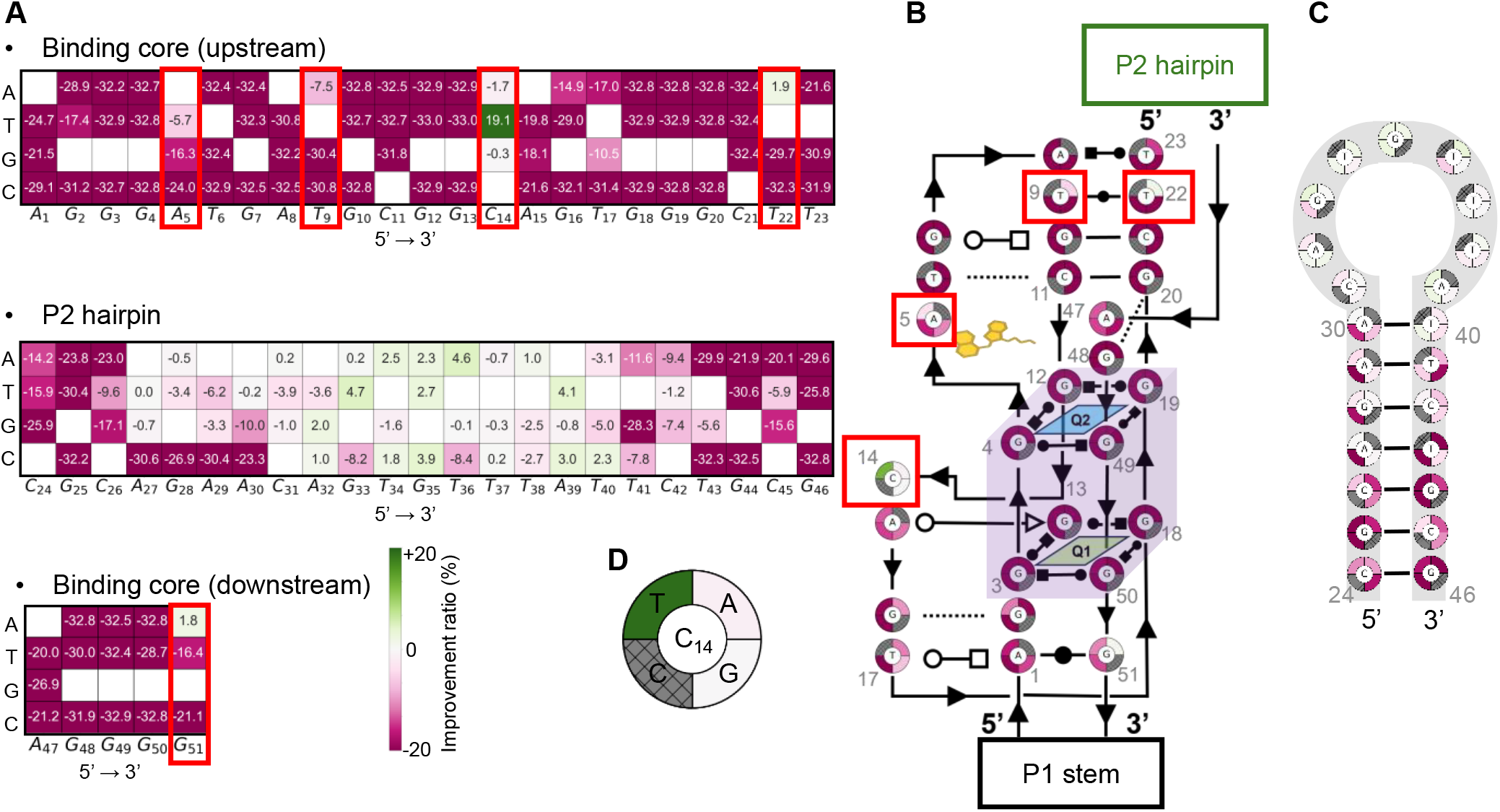
Heatmap of mean improvement ratio for single mutation on NGS chip surface. (**A**) The single mutation heatmap was split into three halves including binding core (upstream), P2 hairpin, and binding core (downstream), in accordance with our library design. The important nucleobases, A5, T9, C14, T22 and G51 were highlighted in red boxes. (**B**) The single mutation heatmap of binding core and (**C**) P2 hairpin. The heatmap was re-organized to show the spatial coordinates based on the secondary structure reported in ref. ^18^. Lines with arrowheads and Leontis–Westhof symbols^43^ denote connectivity and type of base pairs, respectively. In (**C**), A5, T9, C14 and T22 are highlighted in red box due to their unique improvement ratio on the chip surface. Q1 and Q2 represent the two G-quadruplex planes, highlighted in purple, in the canonical Lettuce structure. (**D**) A zoom-in schematic of C14 node. The peripheral four quadrants representing C14A, C14T, C14, C14G from first quadrant to the fourth quadrant, respectively. The quadrant shows the canonical nucleobase are cross-out and label as gray.

To validate the D-FAP/CHAMP screening results, 85 Lettuce variants (out of a library of 8,821 designs) were separately synthesized and measured in test tubes using fluorometry. A good correlation between the CHAMP-derived improvement indices and the fluorometer-based ensemble fluorescence measurements was obtained (R^2^ ∼ 0.78 in **Figure 3A**). As expected, the winner from CHAMP screening, C14T variant, showed an even higher fluorescence improvement in the test tube measurement (45% stronger than the canonical sample as compared to only 19.1% fluorescence increase observed on chip surface). We have previously shown that the test tube validation results typically have the intensity fold change higher than that of the chip screening results, due to the difference in excitation/emission scheme and the high background on chip surface^38^. It has been shown before that changing one nucleobase in an R-FAP (e.g., Broccoli) can alter R-FAP’s emission spectra^39^. Similar emission spectrum shifts were also observed in our D-FAP mutational analysis. Single mutation A5 resulted in a red-shifted emission peak, moving from 558 to 563 nm (**Figure 3B, Table 1**). In contrast, mutations at C14 did not significantly alter the emission peak (**Figure 3C, Table 1**). However, a mutation at another site, A47, caused a blue shift in emission peak from 558 to 552 nm (A47G; **Figure 3D, Table 1**). Interestingly, when the brightest single-mutation variant, C14T, was combined with A47T or A47C, the emission peak seemed to be more influenced by the A47 variants rather than by the C14T (**Figure 3D, Table 1**). This result suggested that while C14T is beneficial in increasing the fluorescence intensity, A47 played a more critical role in determining the emission color.

**Table 1.**
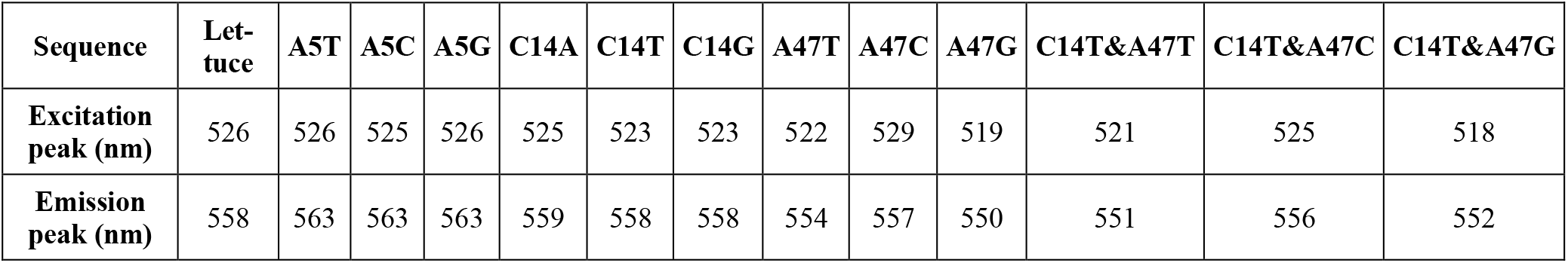
Excitation and emission peaks of selected Lettuce variants.

| Sequence | Lettuce | A5T | A5C | A5G | C14A | C14T | C14G | A47T | A47C | A47G | C14T&A47T | C14T&A47C | C14T&A47G |
| --- | --- | --- | --- | --- | --- | --- | --- | --- | --- | --- | --- | --- | --- |
| Excitation peak (nm) | 526 | 526 | 525 | 526 | 525 | 523 | 523 | 522 | 529 | 519 | 521 | 525 | 518 |
| Emission peak (nm) | 558 | 563 | 563 | 563 | 559 | 558 | 558 | 554 | 557 | 550 | 551 | 556 | 552 |

**Figure 3.**
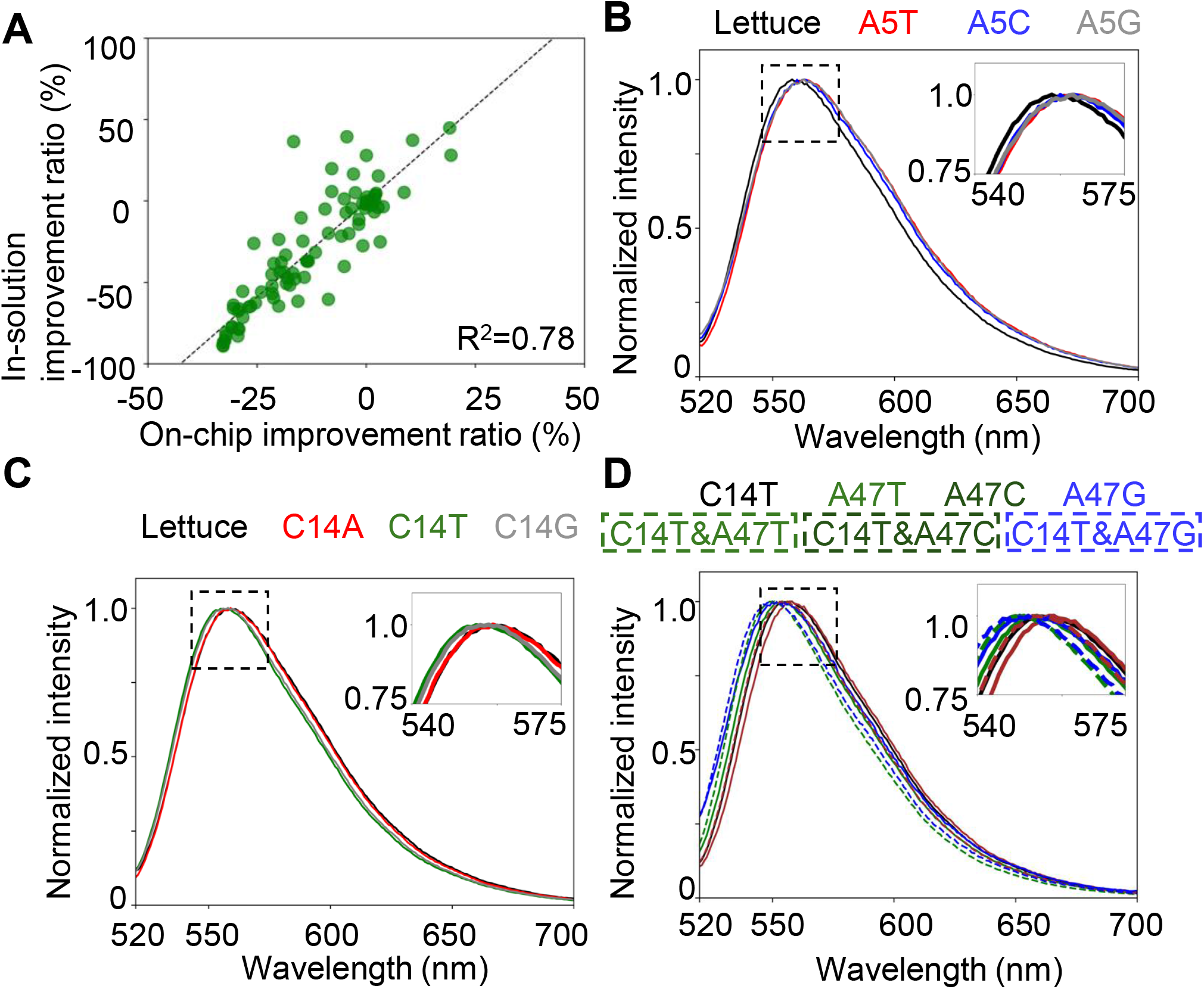
In-solution testing of selected Lettuce variants. (**A**) Randomly selected candidates underwent validation in solution and were subsequently compared to chip results (n = 3). The dotted line represents the linear regression line of the data. The correlation between the in-solution improvement ratio and the on-chip improvement ratio demonstrates a robust representation of chip results in the in-solution outcome. (**B**) The emission spectra of Lettuce, A5T, A5C and A5G. (**C**) The emission spectra of Lettuce, C14A, C14T and C14G. (**D**) The emission spectra of Lettuce, A47T, A47C, A47G, C14T&A47T, C14T&A47C and C14T&A47G. Dotted lines represent double mutation variants. Excitation and emission peaks information are listed in **Table 1**.

The influence of the bases proximal to the binding core on the encapsulated chromophore is not only limited to chromophore’s emission spectrum but also its quantum yield (QY) and fluorescence lifetime. Various fluorescence quantum yields were observed among the selected Lettuce/TO1-biotin variants (QY = 0.59, 0.44, 0.62, 0.44 for canonical Lettuce, A5T, C14T and T22A, respectively; **Supplementary Figure S8)**, whereas the reported QY for Lettuce/DFHBI-1T is only 0.11^17^. In addition, C14T/TO1-biotin showed a fluorescence lifetime (6.00 ns) slightly longer than that of Lettuce/TO1-biotin complex (5.77 ns), while A5T/TO1-biotin gave the shortest fluorescence lifetime (4.83 ns). These lifetime measurement results were consistent with the fluorescence intensity observed both on chip and in test tubes, where C14T and A5T were brighter and dimmer than canonical Lettuce, respectively. Interestingly, we found Lettuce/TO1-biotin and its variants have their apparent dissociation constant 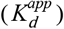 higher than that of Lettuce/DFHBI-1T (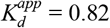, 0.41, 0.63 and 0.61 µM for Lettuce/, A5T/, C14T/ and T22A/TO1-biotin, respectively, and 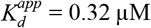 for Lettuce/DFHBI-1T^17^; **Supplementary Figure S7**). This could be explained by the fact that Lettuce was initially selected against DFHBI-1T but not against TO1-biotion. This result also indicated that there could be another D-FAP design that not only binds TO1-biotin strongly 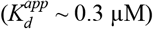 but also shows a high quantum yield (QY > 0.6).

Based on the Lettuce/DFHBI-1T structure, A5 and C14 were the two critical nucleobases that interact with the chromophore via hydrogen bonds and van der Waals (vdW) contacts, respectively^18^. Mutations at A5 position almost entirely diminished Lettuce/DFHBI-1T fluorescence, while C14T only showed a slight decrease in fluorescence^18^. In contrast, our chip screening against TO1-biotin yielded very different results: A5T/TO1-biotin only reduced the fluorescence by 5.7%, while C14T/TO1-biotin even showed 19.1% improvement over the canonical Lettuce/TO1-biotin (**Figure 2A**). From the test-tube validation using fluorometry, the fluorescence decrease of A5T/TO1-biotin changed to 20%, while the fluorescence increase of C14T/TO1-biotin scaled up to 45% over the canonical sample. This suggested that how proximal nucleobases interact with DFHBI-1T and TO1-biotin could be different while the overall crystal structures of Lettuce/ DFHBI-1T and Lettuce/TO1-biotin are similar.

To interrogate the organization of TO1-biotin in Lettuce, we conducted molecular dynamic (MD) simulations on sLettuce, sA5T and sC14T variants, using the reported sLettuce/DFHBI-1T crystal structures (PDB ID: 8FHX and 8FI2^18^) and the AMOEBA^40^ (atomic multipole optimized energetics for biomolecular applications) force fields. System equilibration was assessed by computing the root-mean-square deviations (RMSDs) to the initial structure. While the whole complex was not very stable for some variants (i.e., sC14T), the binding site (defined as all atoms within 10Å of any ligand atom) stayed at minimum fluctuation, indicating that the system has reached equilibrium for binding analysis and that peripheral regions predominantly contribute to the overall RMSDs variance (**Figure 4A-C**). During the sLettuce/TO1-biotin simulation, we observed transient displacement of the potassium ion (K_1_ ^+^) from its pocket. A similar phenomenon was also identified in the simulation of sC14T/TO1-biotin but without permanent displacement (**Figure 4A-C**). Despite this transient displacement, the binding core remained stable, suggesting K_1_ ^+^ is not as critical to the binding of TO1-biotin as to that of DFHBI-1T. We observed no atom in TO1 that could potentially coordinate with K_1_ ^+^, presumably accounting for the tendency of K_1_ ^+^ to move out of place.

**Figure 4.**
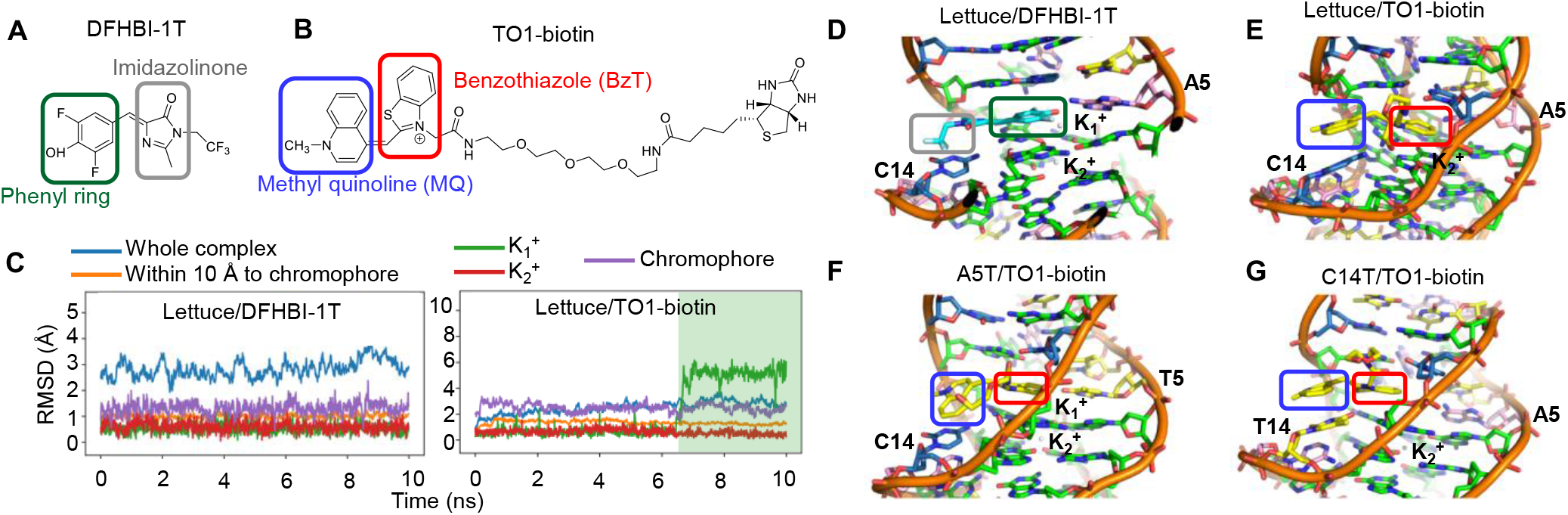
Simulated structures of selected Lettuce variants. (**A**) The structure of DFHBI-1T and (**B**) TO1-biotin. (**C**)We simulated Lettuce/DFHBI-1T, Lettuce/TO1-biotin, A5T/TO1-biotin and C14T/TO1-biotin, using AMEOBA. By computing the structure differences between simulations and crystallography (RMSD), we noticed that one of the potassium ions (i.e., K_1_+) tended to displace from the aptamer core (highlighted in green). (**D-G**) 3D structure of simulated aptamers with chromophores. The structure is based on sequence design of the P1 stem and P2 hairpin reported in ref. ^18^. Moieties are highlighted in colored boxes according to the coloring rules in (**A**) and (**B**).

In comparison to the binding mechanisms of DFHBI-1T to sLettuce (e.g., hydrogen bonds and vdW contacts)^18^, we noticed that, unlike the phenyl ring of DFHBI-1T, the benzothiazole (BzT) group of TO1-biotin lacks the ability to engage neighboring base residues via halogen bonds even though it is still sandwiched in between G12 and C11-G20 base pair. Furthermore, in contrast to the trifluoromethyl group having vdW interactions with C14 in the DFHBI-1T case, we found that the methylquinoline (MQ) group of TO1-biotin establishes π-π stacking. These discoveries indicated that the TO1-biotin interacts with the Lettuce aptamer primarily through π-π stacking rather than hydrogen bonds or vdW contacts (**Figure 4D-G**).

To further assess how stacking interaction could influence the FAP fluorescence, we computed the stacking scores^41^ of the TO1-biotin to the nearby nucleobases in several sLettuce variants (**Supplementary Figure S9**). Intriguingly, we noticed that sC14T/TO1-biotin shows the highest stacking score between T14 and MQ, in consistent with its strongest fluorescence emission (**Supplementary Figure S9C**). In contrast, sA5T mutation led to a lower stacking score compared to that of sLettuce (**Supplementary Figure S9A-B**), which could explain why A5T/TO1-biotin is dimmer than the canonical complex. Furthermore, since T22A formed a T9-A22 base pair on top of the C11-G20 base pair, we hypothesized such a substitution could potentially result in stronger stacking interaction between bases and the BzT group. Indeed, we found an increased stacking score between the BzT group to C11 and G20. However, it is worth noting that the stacking between C14 and MQ is abolished after the T22A mutation (**Supplementary Figure S9D**). Consequently, we concluded that the superior brightness of the Lettuce/TO1-biotin system stems from the rigid stacking interaction of TO1-biotin fluorophore in Lettuce, similar to the scenario in the Mango-II/TO1-biotin complex^20^.

Other thiazole orange chromophores such as TO3-biotin were also developed that show strong binding affinity against existing R-FAPs such as Broccoli and Mango-III^42^. When complexing Lettuce with TO3-biotin, the resulting Lettuce/TO3-biotin sample showed ensemble fluorescence 2.6 times higher than that of Lettuce/DFBHI-1T and a red-shifted emission peak from 505 to 675 nm (**Supplementary figure S2**). Considering the fact that the Lettuce/TO1-biotin’s ensemble fluorescence was 4 times higher than that of Lettuce/DFHBI-1T, here we only focused on Lettuce/TO1-biotin instead of Lettuce/TO3-biotin for mutagenesis studies on NGS chips.

In this study, we discovered a new D-FAP/chromophore system (Lettuce/TO1-biotin) that is 4-fold brighter than its original form (Lettuce/DFHBI-1T) under the same aptamer and chromophore concentrations. This mainly attributes to the increase quantum yield (0.59 vs. 0.11), despite the fact that TO1-biotin has a lower binding affinity to Lettuce as compared to DFHBI-1T (0.82 vs. 0.32 µM). Mutagenesis analysis of 8,821 Lettuce variants on an Illumina NGS chip revealed several interesting mutants that give brighter emission (C14T, 45% brighter than canonical Lettuce), shorter fluorescence lifetime (A5T, 4.83 vs. 5.77 ns), or slightly different emission color (12 nm spectra shift between C14T&A47T and C14T). To explore the chromophore coordination in D-FAP, we simulated the structure of several complexes using AMEOBA force field and found that TO1-biotin primarily bound sLettuce through π-π stacking. The difference in major binding mechanisms between TO1-biotin (π-π stacking) and DFHBI-1T (various types of interactions) might have led to their differences in apparent dissociation constant and quantum yield. In conclusion, our study showcased the feasibility of a massively parallel chip screening platform in optimizing D-FAPs. This platform offered a high-throughput approach to identify potential nucleobases affecting the fluorescence emission without the need of sophisticated instruments and prior knowledge of the aptamer/chromophore complex. We anticipate optimization of more fluorogenic aptamers will soon be carried out using our screening platform.

## ASSOCIATED CONTENT

## AUTHOR INFORMATION

### Corresponding Author

**Hsin-Chih Yeh** – Department of Biomedical Engineering, The University of Texas at Austin, Austin, Texas 78712, USA; Texas Materials Institute, The University of Texas at Austin, Austin, Texas 78712, USA.

### Authors

**Yu-An Kuo** – Department of Biomedical Engineering, The University of Texas at Austin, Austin, Texas 78712, USA

**Yuan-I Chen** – Department of Biomedical Engineering, The University of Texas at Austin, Austin, Texas 78712, USA

**Yanxing Wang** – Department of Biomedical Engineering, The University of Texas at Austin, Austin, Texas 78712, USA

**Zeynep Korkmaz** – Department of Biomedical Engineering, The University of Texas at Austin, Austin, Texas 78712, USA

**Siem Yonas** – Department of Biomedical Engineering, The University of Texas at Austin, Austin, Texas 78712, USA

**Yujie He** – Department of Biomedical Engineering, The University of Texas at Austin, Austin, Texas 78712, USA

**Trung D. Nguyen** – Department of Biomedical Engineering, The University of Texas at Austin, Austin, Texas 78712, USA

**Soonwoo Hong** – Department of Biomedical Engineering, The University of Texas at Austin, Austin, Texas 78712, USA

**Anh Nguyen** – Department of Biomedical Engineering, The University of Texas at Austin, Austin, Texas 78712, USA

**Sohyun Kim** – Department of Biomedical Engineering, The University of Texas at Austin, Austin, Texas 78712, USA

**Saeed Seifi** – Department of Biomedical Engineering, The University of Texas at Austin, Austin, Texas 78712, USA

**Po-Hsun Fan** – Department of Chemistry, The University of Texas at Austin, Austin, Texas 78712, USA

**Yuting Wu** – Department of Chemistry, The University of Texas at Austin, Austin, Texas 78712, USA

**Zhenglin Yang** – Department of Chemistry, The University of Texas at Austin, Austin, Texas 78712, USA

**Hung-Wen Liu** – Department of Chemistry, The University of Texas at Austin, Austin, Texas 78712, USA

**Yi Lu** – Department of Chemistry, The University of Texas at Austin, Austin, Texas 78712, USA

**Pengyu Ren** – Department of Biomedical Engineering, The University of Texas at Austin, Austin, Texas 78712, USA

### Author Contributions

Y.-A. Kuo, Y.-I. Chen, Y. Wang, Y. Wu, Z. Yang, P. Ren, Y. Lu and H.-C. Yeh discussed and defined the project. H.-C. Yeh, P. Ren and Y. Lu supervised the project. Y.-A. Kuo, Y.-I. Chen, Z. Korkmaz and Y. He performed the experiments. H.-W. Liu and Y. Lu provided materials. Y.-A. Kuo, Y.-I. Chen and S. Yonas revised CHAMP algorithms. Y.-A. Kuo, Y.-I. Chen, T. D. Nguyen, S. Hong, A.-T. Nguyen, S. Kim, S. Seifi and P.-H. Fan analyzed the data. Y. Wang and Y. He performed MD simulation. Y.-A. Kuo, Y.-I. Chen, Y. Wang, P. Ren and H.-C. Yeh wrote the article with editorial assistance from all co-authors.

### Competing interests

The authors declare no competing financial interest.

## ACKNOWLEDGMENT

This work was supported by the National Science Foundation grant (CBET2235455 to H.-C. Yeh and Y. Lu) and the National Institutes of Health grant (DA060543 to H.-C. Yeh). P. Ren and Y. Wang are grateful for support by National Institutes of Health (R01GM106137 and R01GM114237), Welch Foundation (F-2120), and the Cancer Prevention and Research Institute of Texas grant (RP210088).

## Notes

### Competing Interest Statement

The authors have declared no competing interest.

## REFERENCES

1. Ellington, A. D.; Szostak, J. W., In vitro selection of RNA molecules that bind specific ligands. Nature 1990, 346 (6287), 818–822.

2. Bock, L. C.; Griffin, L. C.; Latham, J. A.; Vermaas, E. H.; Toole, J. J., Selection of single-stranded DNA molecules that bind and inhibit human thrombin. Nature 1992, 355 (6360), 564–566.

3. Peinetti, A. S.; Lake, R. J.; Cong, W.; Cooper, L.; Wu, Y.; Ma, Y.; Pawel, G. T.; Toimil-Molares, M. E.; Trautmann, C.; Rong, L.; Marinas, B.; Azzaroni, O.; Lu, Y., Direct detection of human adenovirus or SARS-CoV-2 with ability to inform infectivity using DNA aptamer-nanopore sensors. Science Advances 2021, 7 (39), eabh2848.

4. Ellington, A. D.; Szostak, J. W., In vitro selection of RNA molecules that bind specific ligands. Nature 1990, 346 (6287), 818–22.

5. Tuerk, C.; Gold, L., Systematic evolution of ligands by exponential enrichment: RNA ligands to bacteriophage T4 DNA polymerase. Science 1990, 249 (4968), 505–10.

6. Babendure, J. R.; Adams, S. R.; Tsien, R. Y., Aptamers switch on fluorescence of triphenylmethane dyes. J Am Chem Soc 2003, 125 (48), 14716–7.

7. Hou, Q.; Jaffrey, S. R., Synthetic biology tools to promote the folding and function of RNA aptamers in mammalian cells. RNA Biol 2023, 20 (1), 198–206.

8. Lu, X.; Kong, K. Y. S.; Unrau, P. J., Harmonizing the growing fluorogenic RNA aptamer toolbox for RNA detection and imaging. Chem Soc Rev 2023, 52 (12), 4071–4098.

9. Karunanayake Mudiyanselage, A.; Yu, Q.; Leon-Duque, M. A.; Zhao, B.; Wu, R.; You, M., Genetically Encoded Catalytic Hairpin Assembly for Sensitive RNA Imaging in Live Cells. J Am Chem Soc 2018, 140 (28), 8739–8745.

10. Paige, J. S.; Nguyen-Duc, T.; Song, W.; Jaffrey, S. R., Fluorescence imaging of cellular metabolites with RNA. Science 2012, 335 (6073), 1194.

11. You, M.; Litke, J. L.; Jaffrey, S. R., Imaging metabolite dynamics in living cells using a Spinach-based riboswitch. Proc Natl Acad Sci U S A 2015, 112 (21), E2756-65.

12. Kim, H.; Jaffrey, S. R., A Fluorogenic RNA-Based Sensor Activated by Metabolite-Induced RNA Dimerization. Cell Chem Biol 2019, 26 (12), 1725–1731 e6.

13. Litke, J. L.; Jaffrey, S. R., Highly efficient expression of circular RNA aptamers in cells using autocatalytic transcripts. Nat Biotechnol 2019, 37 (6), 667–675.

14. Li, X.; Mo, L.; Litke, J. L.; Dey, S. K.; Suter, S. R.; Jaffrey, S. R., Imaging Intracellular S-Adenosyl Methionine Dynamics in Live Mammalian Cells with a Genetically Encoded Red Fluorescent RNA-Based Sensor. J Am Chem Soc 2020, 142 (33), 14117–14124.

15. Moon, J. D.; Wu, J.; Dey, S. K.; Litke, J. L.; Li, X.; Kim, H.; Jaffrey, S. R., Naturally occurring three-way junctions can be repurposed as genetically encoded RNA-based sensors. Cell Chem Biol 2021, 28 (11), 1569–1580 e4.

16. Dey, S. K.; Filonov, G. S.; Olarerin-George, A. O.; Jackson, B. T.; Finley, L. W. S.; Jaffrey, S. R., Repurposing an adenine riboswitch into a fluorogenic imaging and sensing tag. Nat Chem Biol 2022, 18 (2), 180–190.

17. VarnBuhler, B. S.; Moon, J.; Dey, S. K.; Wu, J.; Jaffrey, S. R., Detection of SARS-CoV-2 RNA Using a DNA Aptamer Mimic of Green Fluorescent Protein. ACS Chem Biol 2022, 17 (4), 840–853.

18. Passalacqua, L. F. M.; Banco, M. T.; Moon, J. D.; Li, X.; Jaffrey, S. R.; Ferre-D’Amare, A. R., Intricate 3D architecture of a DNA mimic of GFP. Nature 2023, 618 (7967), 1078–1084.

19. Trachman, R. J., 3rd; Autour, A.; Jeng, S. C. Y.; Abdolahzadeh, A.; Andreoni, A.; Cojocaru, R.; Garipov, R.; Dolgosheina, E. V.; Knutson, J. R.; Ryckelynck, M.; Unrau, P. J.; Ferre-D’Amare, A. R., Structure and functional reselection of the Mango-III fluorogenic RNA aptamer. Nat Chem Biol 2019, 15 (5), 472–479.

20. Trachman, R. J., 3rd; Abdolahzadeh, A.; Andreoni, A.; Cojocaru, R.; Knutson, J. R.; Ryckelynck, M.; Unrau, P. J.; Ferre-D’Amare, A. R., Crystal Structures of the Mango-II RNA Aptamer Reveal Heterogeneous Fluorophore Binding and Guide Engineering of Variants with Improved Selectivity and Brightness. Biochemistry 2018, 57 (26), 3544–3548.

21. Buenrostro, J. D.; Araya, C. L.; Chircus, L. M.; Layton, C. J.; Chang, H. Y.; Snyder, M. P.; Greenleaf, W. J., Quantitative analysis of RNA-protein interactions on a massively parallel array reveals biophysical and evolutionary landscapes. Nat Biotechnol 2014, 32 (6), 562–8.

22. Boyle, E. A.; Andreasson, J. O. L.; Chircus, L. M.; Sternberg, S. H.; Wu, M. J.; Guegler, C. K.; Doudna, J. A.; Greenleaf, W. J., High-throughput biochemical profiling reveals sequence determinants of dCas9 off-target binding and unbinding. Proc Natl Acad Sci U S A 2017, 114 (21), 5461–5466.

23. Denny, S. K.; Bisaria, N.; Yesselman, J. D.; Das, R.; Herschlag, D.; Greenleaf, W. J., High-Throughput Investigation of Diverse Junction Elements in RNA Tertiary Folding. Cell 2018, 174 (2), 377–390 e20.

24. Layton, C. J.; McMahon, P. L.; Greenleaf, W. J., Large-Scale, Quantitative Protein Assays on a High-Throughput DNA Sequencing Chip. Mol Cell 2019, 73 (5), 1075–1082 e4.

25. Andreasson, J. O. L.; Savinov, A.; Block, S. M.; Greenleaf, W. J., Comprehensive sequence-to-function mapping of cofactor-dependent RNA catalysis in the glmS ribozyme. Nat Commun 2020, 11 (1), 1663.

26. Jung, C.; Hawkins, J. A.; Jones, S. K., Jr.; Xiao, Y.; Rybarski, J. R.; Dillard, K. E.; Hussmann, J.; Saifuddin, F. A.; Savran, C. A.; Ellington, A. D.; Ke, A.; Press, W. H.; Finkelstein, I. J., Massively Parallel Biophysical Analysis of CRISPR-Cas Complexes on Next Generation Sequencing Chips. Cell 2017, 170 (1), 35–47.

27. Jones, S. K., Jr.; Hawkins, J. A.; Johnson, N. V.; Jung, C.; Hu, K.; Rybarski, J. R.; Chen, J. S.; Doudna, J. A.; Press, W. H.; Finkelstein, I. J., Massively parallel kinetic profiling of natural and engineered CRISPR nucleases. Nat Biotechnol 2021, 39 (1), 84–93.

28. Kuo, H. C.; Prupes, J.; Chou, C. W.; Finkelstein, I. J., Massively Parallel Profiling of RNA-targeting CRISPR-Cas13d. bioRxiv 2023.

29. Wu, D.; Feagin, T.; Mage, P.; Rangel, A.; Wan, L.; Kong, D.; Li, A.; Coller, J.; Eisenstein, M.; Soh, H., Flow-Cell-Based Technology for Massively Parallel Characterization of Base-Modified DNA Aptamers. Anal Chem 2023, 95 (5), 2645–2652.

30. Yoshikawa, A. M.; Rangel, A. E.; Zheng, L.; Wan, L.; Hein, L. A.; Hariri, A. A.; Eisenstein, M.; Soh, H. T., A massively parallel screening platform for converting aptamers into molecular switches. Nat Commun 2023, 14 (1), 2336.

31. Wu, D.; Feagin, T.; Mage, P.; Rangel, A.; Wan, L.; Kong, D.; Li, A.; Coller, J.; Eisenstein, M.; Soh, H. T., Automated platform for high-throughput screening of base-modified aptamers for affinity and specificity. bioRxiv 2022.

32. Kuo, Y. A.; Jung, C.; Chen, Y. A.; Kuo, H. C.; Zhao, O. S.; Nguyen, T. D.; Rybarski, J. R.; Hong, S.; Chen, Y. I.; Wylie, D. C.; Hawkins, J. A.; Walker, J. N.; Shields, S. W. J.; Brodbelt, J. S.; Petty, J. T.; Finkelstein, I. J.; Yeh, H. C., Massively Parallel Selection of NanoCluster Beacons. Adv Mater 2022, e2204957.

33. Nutiu, R.; Friedman, R. C.; Luo, S.; Khrebtukova, I.; Silva, D.; Li, R.; Zhang, L.; Schroth, G. P.; Burge, C. B., Direct measurement of DNA affinity landscapes on a high-throughput sequencing instrument. Nat Biotechnol 2011, 29 (7), 659–64.

34. Tome, J. M.; Ozer, A.; Pagano, J. M.; Gheba, D.; Schroth, G. P.; Lis, J. T., Comprehensive analysis of RNA-protein interactions by high-throughput sequencing-RNA affinity profiling. Nat Methods 2014, 11 (6), 683–8.

35. Svensen, N.; Peersen, O. B.; Jaffrey, S. R., Peptide Synthesis on a Next-Generation DNA Sequencing Platform. Chembiochem 2016, 17 (17), 1628–35.

36. Dolgosheina, E. V.; Jeng, S. C.; Panchapakesan, S. S.; Cojocaru, R.; Chen, P. S.; Wilson, P. D.; Hawkins, N.; Wiggins, P. A.; Unrau, P. J., RNA mango aptamer-fluorophore: a bright, high-affinity complex for RNA labeling and tracking. ACS Chem Biol 2014, 9 (10), 2412–20.

37. Autour, A.; S, C. Y. J.; A. D. C.; Abdolahzadeh, A.; Galli, A.; Panchapakesan, S. S. S.; Rueda, D.; Ryckelynck, M.; Unrau, P. J., Fluorogenic RNA Mango aptamers for imaging small non-coding RNAs in mammalian cells. Nat Commun 2018, 9 (1), 656.

38. Kuo, Y.-A.; Jung, C.; Chen, Y.-A.; Kuo, H.-C.; Zhao, O. S.; Nguyen, T. D.; Rybarski, J. R.; Hong, S.; Chen, Y.-I.; Wylie, D. C.; Hawkins, J. A.; Walker, N. W.; Shields, S. W. J.; Brodbelt, J. S.; Petty, J. T.; Finkelstein, I. J.; Yeh, H.-C., Massively parallel selection of NanoCluster Beacons. Adv Mater 2022, 2204957.

39. Filonov, G. S.; Song, W.; Jaffrey, S. R., Spectral Tuning by a Single Nucleotide Controls the Fluorescence Properties of a Fluorogenic Aptamer. Biochemistry 2019, 58 (12), 1560–1564.

40. Yang, X.; Liu, C.; Kuo, Y. A.; Yeh, H. C.; Ren, P., Computational study on the binding of Mango-II RNA aptamer and fluorogen using the polarizable force field AMOEBA. Front Mol Biosci 2022, 9, 946708.

41. Taghavi, A.; Riveros, I.; Wales, D. J.; Yildirim, I., Evaluating Geometric Definitions of Stacking for RNA Dinucleoside Monophosphates Using Molecular Mechanics Calculations. J Chem Theory Comput 2022, 18 (6), 3637–3653.

42. Jeng, S. C. Y.; Trachman, R. J., 3rd; Weissenboeck, F.; Truong, L.; Link, K. A.; Jepsen, M. D. E.; Knutson, J. R.; Andersen, E. S.; Ferre-D’Amare, A. R.; Unrau, P. J., Fluorogenic aptamers resolve the flexibility of RNA junctions using orientation-dependent FRET. RNA 2021, 27 (4), 433–444.

43. Leontis, N. B.; Westhof, E., Geometric nomenclature and classification of RNA base pairs. RNA 2001, 7 (4), 499–512.

